# Single-cell RNA sequencing reveals recruitment of the M2-like CCL8^high^ macrophages in Lewis lung carcinoma-bearing mice following hypofractionated radiotherapy

**DOI:** 10.1101/2024.01.15.575792

**Authors:** Haonan Yang, Zheng Lei, Jiang He, Lu Zhang, Tangmin Lai, Liu Zhou, Nuohan Wang, Zheng Tang, Jiangdong Sui, Yongzhong Wu

## Abstract

Tumor-associated macrophages (TAMs) are a crucial factor in reprogramming the tumor microenvironment following radiotherapy. The mechanisms underlying this process remain to be elucidated. Here, we seek to investigate the effects of hypofractionated radiotherapy on macrophages dynamics in a subcutaneous Lewis lung carcinoma murine model. Utilizing single-cell RNA sequencing, we identified a distinct M2-like population of macrophages with high Ccl8 expression level post-hypofractionated radiotherapy. Remarkbly, hypofractionated radiotherapy promoted CCL8^high^ macrophages infiltration and reprogrammed them by upregulating immunosuppressive genes and downregulating antigen-presenting genes, leading to an immunosuppressive tumor microenvironment. Further, we demonstrated that hypofractionated radiotherapy amplified the CCL signaling pathway, enhancing the pro-tumorigenic functions of CCL8^high^ macrophages and promoting macrophages recruitment. The combination therapy of hypofractionated radiotherapy with the CCL signal inhibitor Bindarit was effective in reducing M2 macrophages infiltration and extending the duration of local tumor control. This research highlights the potential of targeting TAMs and introduces a novel combination to improve the efficacy of hypofractionated radiotherapy.

## Introduction

Radiation therapy is an important modality for cancer treatment. In the case of non-small cell lung cancer (NSCLC), radiotherapy is the only treatment applicable to all TNM stages[1]. It is estimated that over 60% of NSCLC patients require radiotherapy during their treatment[2]. Hypofractionated radiotherapy, especially for the Stereotactic body radiation therapy (SBRT), is a precision treatment technique that targets the tumor while minimizing exposure to surrounding normal tissue. This allows for high-dose fractionated radiation, shortening treatment duration and improving therapeutic outcomes. Specifically, for inoperable early-stage NSCLC, hypofractionated radiotherapy significantly improves clinical outcomes compared to conventional radiotherapy and has become the current standard treatment protocol[3,4]. Beyond its direct tumoricidal effects, hypofractionated radiotherapy also modulates immune response through various mechanisms including the release of tumor-associated antigens (neoantigens) and damage-associated molecular patterns (DAMPs), enhancement of antigen presentation, releasing of cytokines and chemokines for inflammatory response regulation, and promotion of T cell activation and infiltration[5–8]. However, hypofractionated radiotherapy also contributes to the recruitment of immunosuppressive cells such as FOXP3^+^ regulatory T cells (Tregs), tumor-associated neutrophils (TANs), tumor-associated macrophages (TAMS), and myeloid-derived suppressor cells (MDSCs) via a series of cytokines and chemokines, thereby inducing an immunosuppressive tumor microenvironment[9,10].

Tumor-associated macrophages are traditionally categorized into two phenotypes: M1 and M2. M1 macrophages are considered pro-inflammatory and play a role in inhibiting tumor growth, whereas M2 macrophages exhibit anti-inflammatory functions and are known to promote tumor progression[11,12]. M2 TAMs form an important component of tumor immunosuppressive microenvironment[13,14]. Radiotherapy has been shown to promote the infiltration of TAMs into tumors which can attenuate the therapeutic efficacy of radiotherapy[5]. As a result, understanding the dynamics of TAMs in the context of radiotherapy and targeting TAMS to overcome this immunosuppressive effect is vital for enhancing treatment efficacy.

Chemokines are a group of cytokines that induce cell migration and participate in regulating inflammatory responses by binding to corresponding receptors. Based on the N-terminal cysteine residue sequence, chemokines are classified into four subtypes: CC, CXC, C, and CX3C, among which CC chemokines are further divided into 27 types14. In the tumor microenvironment, CC chemokines mediate the infiltration and differentiation of immunosuppressive cells such as TAM, MDSC, TAN, Treg, and contribute to tumor immune evasion[15,16]. Current research finds that radiotherapy recruits immunosuppressive cells to reshape the tumor microenvironment, weakening anti-tumor immunity, with CC chemokines playing a significant role in this process[17]. Specifically, CCL2 (C-C motif ligand 2) is most clearly related to the post-radiotherapy immune microenvironment. It not only recruits immunosuppressive cells such as tumor-associated macrophages, myeloid-derived suppressor cells (MDSC) but also induces macrophage polarization and further secretion of cytokines[18,19]. Unlike CCL2, tumor cell release CCL5 promotes TAM and MDSC infiltration, participating in tumor metastasis[20]; it also recruits CD8+ T cells enhancing adaptive immunity, demonstrating dual immune regulatory roles[21]. CCL7 is reported to polarize macrophage as M1 type and mediating radiation-induced lung injury[22]. The role of CCL8 in reshaping the tumor microenvironment following radiotherapy is not yet clear, but it is known to promote tumor cell proliferation and migration abilities[23,24].

In the present study, we profiled early alterations of tumor microenvironment in a subcutaneous (s.c.) Lewis lung carcinoma (LLC) murine model following hypofractionated radiotherapy by performing single-cell RNA sequencing. We then identified a M2-like CCL8^high^ macrophage population which contributed to immune suppression. Moreover, hypofractionated radiotherapy promoted this specific macrophage population infiltration and reprogrammed TAMs through CCL signaling pathway. The CCL signals inhibitor, Bindarit, synergized with hypofractionated radiotherapy to extend local control in our s.c. LLC murine model.

## Methods

### Cell lines and drugs

Lewis lung carcinoma (LLC) cell line was purchased from the National Collection of Authenticated Cell Cultures, China. Cells were cultured at 37 ° humidified incubator with 5% CO2 in DMEM medium supplemented with 10% FBS and 100 units/ml penicillin and 0.1mg/ml streptomycin. The cells were routinely tested to confirm the absence of Mycoplasma contamination and were cultured for a limited number of generations. The CCL8 inhibitor Bindarit (AF2838, MCE, Cat#: HY-B0498) was diluted in 10% DMSO, then 45% PEG300 (MCE, Cat#: HY-Y0873) and 45% Saline. Recombinant CCL8 protein was purchased from MCE (Cat. #: HY-P7771).

### Mice, in vivo studies, and treatments

Female C57BL/6N mice ages 6 to 8 weeks were purchased from Beijing Vital River Laboratory Animal Technology. All animal procedures followed the Guide for the Care and Use of Laboratory Animals. For the LLC xenograft tumor models, 1 x 10^6^ LLC cells in 100ul PBS were injected subcutaneously in the right flank of C57BL/6 mice. For radiotherapy, once tumors reached an average volume of 100mm^3^, radiation was given on day 7-9 post-tumor cell injection with total dose 24Gy in 3 fractions (8Gy each fraction). For CC chemokines inhibitor, Bindarit was administered intraperitoneally 100mg/kg daily from day 5 post-tumor injection and was continued for 7 days. Tumors were measured with a caliper and mice were euthanized when tumor volumes reached 1500 mm^3^. Weight was monitored three times a week.

### Cell preparation and single-cell RNA sequencing

Tumors were dissociated using Multi Tissue Dissociation Kit 2 (Miltenyi Biotec, Cat#: 130-110-203) according to manufacturer’s protocol. Debris and dead cells were removed (Miltenyi Cat#: 130-109-398/130-090-101). Fresh cells were resuspended at 1×10^6^ cells per ml in 1×PBS and 0.04% bovine serum albumin. Single-cell RNA-Seq libraries were prepared using SeekOne MM Single Cell 3’ library preparation kit following the manufacturer’s instructions. Sequencing was performed on the Illumina NovaSeq 6000 with PE150 read length. The SeekOne Tools pipeline was used to process the cleaned reads and generated the transcript expression matrices. The schematic workflow for single-cell RNA sequencing was generated by using the Biorender software (www.biorender.com).

### Clustering scRNA-seq data and cell type annotation

All additional analyses were performed using R 4.3.0. Unsupervised clustering was performed by Seurat package[25] (version 4.3.0) and cells were integrated using the Harmony[26]. After that highly variable genes (HVGs) were selected for principle components analysis (PCA), and the top 30 significant principal components (PCs) were selected for Uniform Manifold Approximation and Projection (UMAP) and t-distributed stochastic neighbor embedding (tSNE) dimension reduction, and visualization of gene expression. Next, differentially expressed genes (DEGs) of each cell subcluster were identified by running the “FindAllMarkers” function of Seurat package. Final, cell types were annotated according to the expression level of the known canonical marker genes of certain cell types. Transcription factor activity analysis based on DEGs was conducted by using the DoRothEA regulatory network analysis[27].

### Enrichment analysis

DEGs which |logFC| > 0.5 and adjusted P value < 0.05 were selected for further analysis. The clusterProfiler package[28] was used to process GO and KEGG enrichment between macrophage subclusters with or without treatment. Gene set variation analysis of hallmark gene sets (MSigDB) for tumor subclusters was conducted using the GSVA[29].

### Pseudotime analysis

To analyze the gene characteristics of myeloid cells differentiation, pseudotime analysis was performed by monocle2 package[30]. The dpFeature method provided by the differentialGeneTest function was applied to find DEGs for pseudotime analysis. The function called plot_genes_in_pseudotime and plot_pseudotime_heatmap were applied for visualization.

### Cellchat analysis

To explore cell-cell communication between tumor cells and myeloid cells, the Cellchat package[31] was first used to calculate the communication probability of control and IR group separately. Then comparison analysis was applied with the mergeCellChat function to identify the upregulated and downregulated signals in IR group.

### TCGA data analysis

The TCGAbiolinks package was used to download the expression data of TCGA. For the gene signature, we used the three marker genes Ccl8, C1qb, and Fth1. The survival analysis was conducted in accordance with the methods reported by Cheng et al. in their study[32].

### Flow cytometry analysis

Tumors were collected 3 days after last radiation fraction then were dissociated in DMEM supplemented with 1mg/ml Collagenase IV(Solarbio) and 1ul/ml DNase I (Solarbio) at 37 ° for one hour. Single-cell suspensions were collected after filtering digested tissues with 40-um filter and washed with PBS supplemented with 2% FBS. Subsequently, cell surface and intracellular markers were stained with the following fluorochrome-conjugated antibodies: anti-CD45 BV421 (Cat#: 103134), anti-CD11b PE (Cat#: 101207), anti-F4/80 AF700 (Cat#: 123129), anti-CD86 FITC (Cat#: 105109), anti-CD206 APC (Cat#: 141708), anti-CD4 BV510 (Cat#: 100553), anti-CD8a AF700 (Cat#: 100729) from Biolegend. For surface staining, all samples were stained with antibodies at 4° for 30 minutes; for intracellular staining, cells were fixed and permeabilized with BD Cytofix/Cytoperm (Cat#: 554714) then stained according to the manufacturer’s instructions. Analysis of stained cells was performed using a CytoFlex cytometer (Beckman Coulter) and CytExpert software.

### Cytokine and Chemokine assays

Blood of mice bearing LLC tumors were collected 3 days after last radiation fraction. Serum was diluted five times in PBS. Cytokines were quantified according to the manufacturer’s protocol (Luminex Mouse Discovery Assay, R&D Systems). CSF1, CCL2, CCL7, and CCL8 were further quantified by Elisa analysis according to manufacturer’s protocol (MEIMIAN, Cat#: MM-1025M1, MM-0082M1, MM-0084M1, and MM-0083M1).

### Real-Time qPCR

Cells were kept frozen (−80°) until mRNA extraction. The RNeasy kit and the cDNA Synthesis Kit (Takara, Cat#: 9767, RR047A) was used to extract total RNA and prepare cDNA following the manufacturer’s instructions. Real-Time qPCR was performed with BioRad PCR system. Primers for mouse Csf1 were 5’-TAGAAAGGATTCTATGCTGGG-3’ and reverse 5’-CTCTTTGGTTGAGAGTCTAAG-3’ (PrimerBank). They were purchased from Sangon Biotech (Shanghai, China).

### Immunostaining

Tissues were fixed with 4% paraformaldehyde for 72 hours at 4° before being embedded. Immunostaining was performed using the multiplex immunohistochemistry kit from the AiFang biological (China, catalog Cat#: AFIHC035). The following primary antibodies were used: rabbit anti-F4/80 (AiFang biological, Cat#: SAF002), rabbit anti-CD206 (AiFang biological, Cat#: AF07082), rat anti-CCL8 (R&D, Cat#: MAB790). Nuclear staining was performed with DAPI. Images from the stained slides were scanned using the Digital Pathology Slide Scanner (KFBIO, China).

### Cell proliferation assay and Colony formation assay

For proliferation assay and colony formation assay, the LLC cell line was treated with recombinant CCL8 (MedChemExpress, Cat#: HY-P7771) at 0, 5ng/ml, 20ng/ml. Then proliferation was assessed with a Cell Counting Kit-8 (Sigma-Aldrich, Cat#: 96992) following manufacturer’s protocol. For colony formation assay, LLC cells were seeded into 6-well plates. The cell density for each cell was 500. After treating with recombinant CCL8 for 48 hours, the culture medium was replaced by fresh DMEM. Cells were further incubated for another 7 days.

### Statistical analysis

Statistical analyses were conducted by R version 4.3.0 and GranphPad Prism 8.0. Multiple comparisons such as tumor growth curves were analyzed by two-way ANOVA. Violin plots comparing the gene expression level between two groups were analyzed by unpaired two-sided Wilcoxon test. Bar charts with mean value were analyzed by student’s t tests. P values less than 0.05 were considered statistically significant.

## Results

### Single-cell transcriptomic landscape of subcutaneous LLC tumors at early-stage following hypofractionated radiotherapy

To investigate early response of the tumor microenvironment (TME) in lung cancer treated with hypofractionated radiotherapy, single-cell RNA sequencing (scRNA-seq) was performed to 4 subcutaneous (s.c.) LLC tumors at 72 hours following radiation (Figure 1A). Applying established quality control methods, 54883 cells were obtained, including 44919 cells from the hypofractionated radiotherapy group (3 individual samples) and 9964 cells from control mice(3 samples mixed in 1)[6]. All cells were divided into 21 clusters by running the Seurat pipeline, then we identified 8 populations, including tumors cells, macrophages, monocytes, T cells, neutrophils, fibroblasts, endothelial cells, and dendritic cells by applying canonical gene signatures and SingleR package (Figure 1B, D). Marker genes of each cell type were shown by a dot plot (Figure 1 E). Tumor cells were further confirmed by applying the inferCNV analysis. Comparing with macrophages and monocytes (reference cells), tumor cells showed obviously higher frequency of copy number variation (Supplementary Figure S1C). As expected, the ratio of tumor cells decreased after hypofractionated radiotherapy and T cells infiltration was minimal in two groups. Intriguingly, macrophages were highly enriched in hypofractionated radiotherapy treated s.c. LLC tumors (22.41% for RT, 10.19% for NT) (Figure 1C).

**Figure 1:**
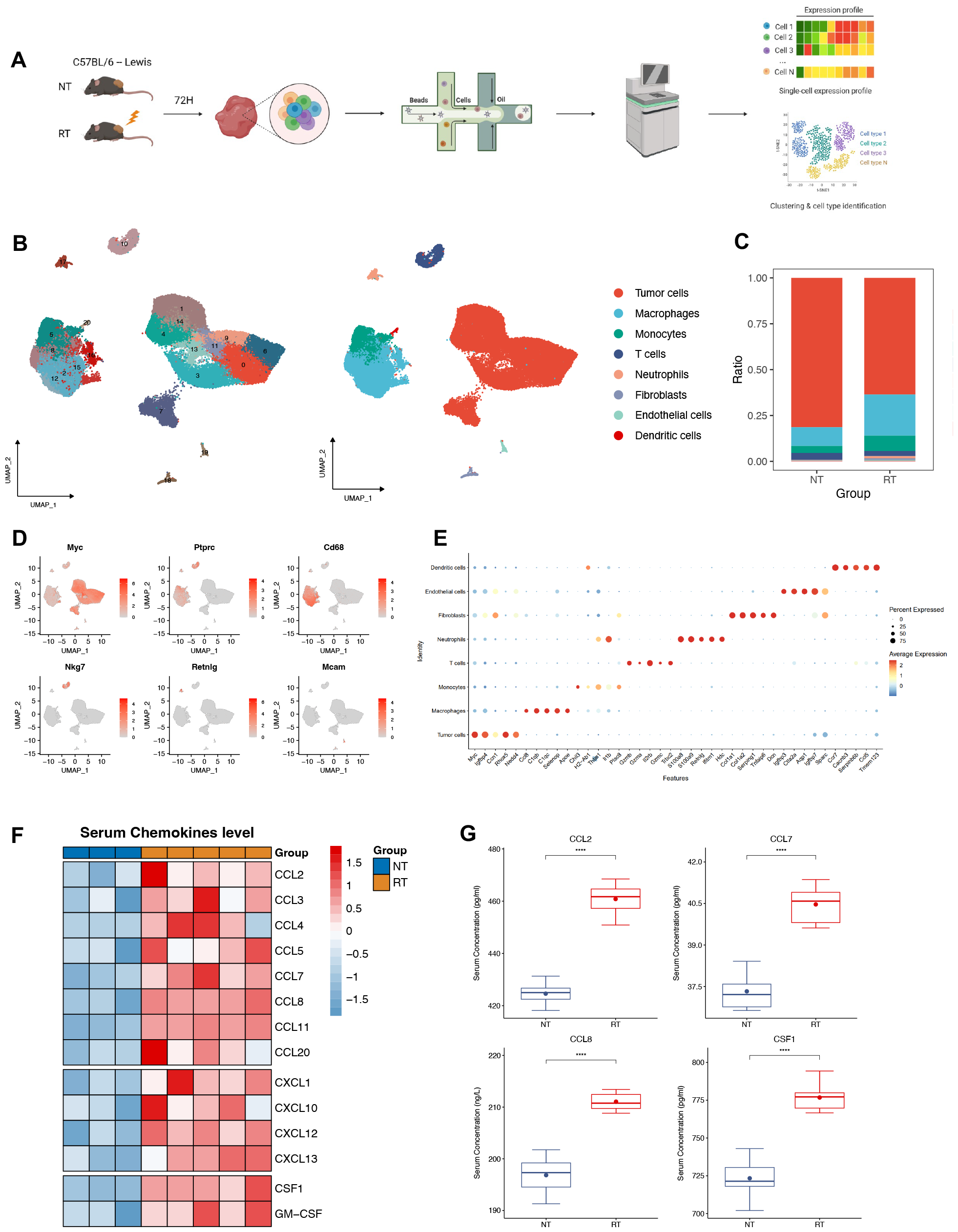
Single-cell transcriptomic landscape of subcutaneous LLC tumors at early-stage following hypofractionated radiotherapy. (A) Overview of the experimental design for single-cell RNA sequencing. (B) Uniform Manifold Approximation (UMAP) plot showing the unsupervised clusters of 54883 single cells and annotated cell types. (C) The proportion of each cell types in LLC tumors treated with or without radiation. For (B and C), each color represents the same cell type. (D) Feature plot and (E) Dot plot showing marker genes. (F) Heatmap showing serum chemokines level in LLC murine models 72 hours post-treatment. Each row in the heatmap has been scaled. (G) The concentrations of CCL2, CCL7, CCL8, and CSF1 in serum measured by ELISA.

To further explore the early impact of hypofractionated radiotherapy on myeloid cells recruitment-related chemokines in lung cancer, multi-chemokines analysis was conducted to evaluate the release of serum chemokines of s.c. LLC bearing mice at 72 hours after hypofractionated radiotherapy. We found that radiation (RT) at subcutaneous tumor site triggered the release of C-C and CXC chemokines at early-stage compared to radiation naive group (NT) (Figure 1F). Consistently, a markedly increased CCL2, CCL7, and CCL8 serum levels after radiation were further determined using the enzyme-linked immunosorbent assay (ELISA) kits (Figure 1G).

### The alteration of LLC cell states following hypofractionated radiotherapy

Performing unsupervised dimensionality reduction and clustering to previously annotated tumor cells, we identified 10 tumor cell populations with distinct gene expression profiles (Figure 2A). The distribution of tumor cell populations was identical in two groups after batch effect correction (Figure 2B). However, cluster 0 and cluster 3 to 5 were less abundant in RT group, while cluster 1, 2, 6 and 7 were enriched after RT (Figure 2C). We then evaluated the stemness of each cell population by calculating the expression score of a curated stemness gene signature[33]. Cluster 0 and 3 demonstrated the highest stemness score among all LLC cells (Figure 2D). These findings were further corroborated using CytoTRACE, a computational algorithm that predict cellular differentiation state, with higher score indicating greater stemness[34]. Based on the differentiation score, cluster 0 and 3 exhibited highest level of stemness (Figure 2E, F), whereas cluster 2 and 8 were identified as the most differentiation tumor cells. Gene set variation analysis (GSVA) revealed that cluster 0 were enriched for expression genes related with DNA repair, cell cycle (Hallmarks E2F-Targets, G2M checkpoint), MYC targets, and Oxidative phosphorylation, which implied higher intrinsic radiation sensitivity. Along with enrichment of DNA repair and cell cycle genes, cluster 3 exhibited higher expression level of epithelial-mesenchymal transition (EMT) related genes. Conversely, cluster 2 and 8 downregulated most hallmark gene sets compared to other clusters (Figure 2 G). Further gene expression profiling analysis confirmed the upregulation of cell cycle genes of cluster 0 and 3. Intriguingly, cluster 6, which exhibited moderate differentiation state, was found highest expression level of transcription factors STAT1/STAT2-IRF9/IRF1 activated by interferon gamma (IFN-γ) leading to upregulation of tumor PD-L1 expression following RT (Fig. 2H). Cell-cell communication analysis between tumor cells and immune cells revealed a specific CSF (colony stimulating factor) signaling pathway recruiting monocytes and macrophages since only tumor cells expressing the ligand gene Csf1 and barely expressing Csf1 receptor gene (Figure 2 I, J and Figure S1D). Moreover, radiation augmented expression of Csf1 in LLC cells and its release in mouse serum (Figure 2 K and Figure 1 G). This was further demonstrated by reverse transcription quantitative PCR(RT-qPCR) (Figure 2 L). These results suggest that LLC maintains heterogeneity in murine s.c. tumor model and enables us to understand the unique transcriptomic characteristics of subpopulations of LLC cells following radiation. Notably, hypofractionated radiotherapy observably promotes CSF signaling pathway, which is supposed to mediate myeloid cells development[35].

**Figure 2:**
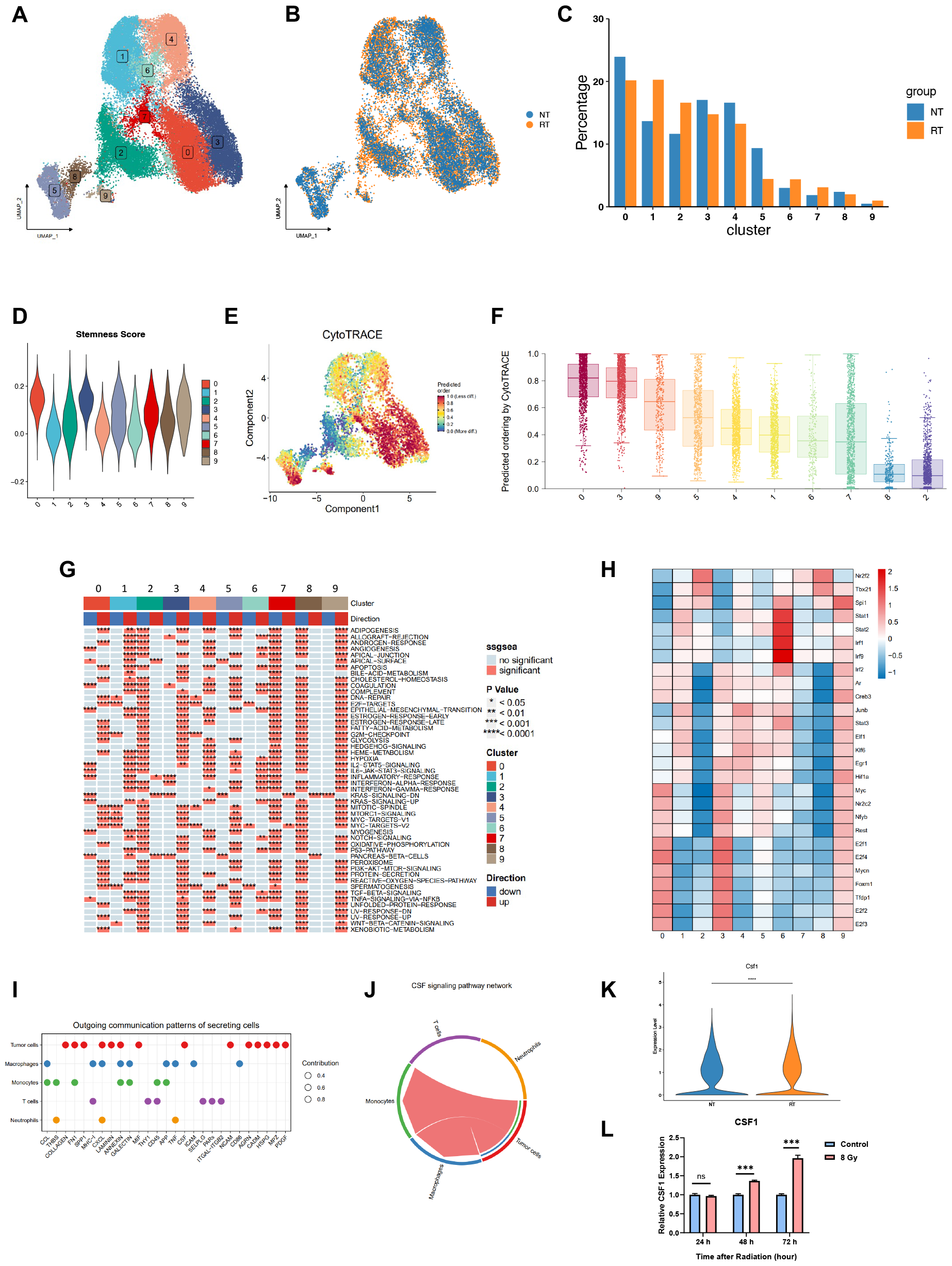
The alteration of LLC cell states following hypofractionated radiotherapy. (A) UMAP plot showing clusters of LLC cells. (B) UMAP plot showing the distribution of LLC cells in each group. (C) The proportion of each cluster in control (NT) and radiotherapy (RT) groups. (D) Violin plot showing stemness score for each cluster. (E) UMAP plot and (F) Bar plot of stemness in LLC cells predicted by CytoTRACE. (G) Heatmap of hallmark gene sets from the MSigDB enriched in different types of cell clusters. (H) Heatmap of transcription factors activity in each cell cluster. (I) Dot plot showing outgoing communication signaling pathways of different cell types. (J) Chord plot of communication network of CCL signaling pathway between tumor cells and other cell types. (K) Csf1 gene expression level between NT and RT groups. (L) The relative mRNA expression levels of Csf1 in LLC cells with or without exposure to 8 Gy of radiation in vitro.

### Hypofractionated radiotherapy stimulates a distinctive immune microenvironment at early-stage

To investigate the influence of hypofractionated radiotherapy on myeloid cells recruitment at early-stage in s.c. LLC tumors, unsupervised clustering and manual annotation identified 6 macrophages clusters (Mac_Ccl8, Mac_Spp1, Mac_Igfbp4, Mac_Hmox1, Mac_Ftl1, and Mac_Stmn1), 2 monocytes clusters (Mono_Plac8 and Mono_Cxcl3) and a cluster of DCs with distinct gene expression pattern (Figure 3 A, C). Marker gene feature plots further validated cell type annotation (Figure 3B). Apoe, representing lipid-associated macrophage or TREM2 macrophage, was mainly expressed in Ccl8^high^ macrophage (Mac_Ccl8), while monocyte or monocyte-derived cells gene, such as Thbs1, was expressed in Mono_Plac8, Mono_Cxcl3 and Mac_Spp1. Notably, Mac_Ccl8 was the major population of tumor-infiltrating myeloid cells, and hypofractionated radiotherapy induced enrichment of Mac_Ccl8 compared to NT (32.28% in RT versus 23.48% in NT) (Figure 3D). Using M1/M2 macrophages gene signatures[32], we found M2 was the dominant polarization state of myeloid cells in LLC-bearing mice. Moreover, Mac_Ccl8 not only exhibited highest M2 signatures score among all TAMs populations but also expressed higher phagocytosis genes relative to other populations (Figure 3E). In addition, the significant correlation of Ccl8 and M2 macrophages was then demonstrated by applying TIMER 2.0 based on TCGA-LUAD expression profile (Figure 3F). Consistently, the Gene Ontology (GO) enrichment analysis indicated upregulation of the chemokine signaling pathway, cytokine-cytokine receptor interaction, antigen processing and presentation, and phagosome gene sets of Mac_Ccl8 (Figure 3G), indicating its multifarious functional features. When compared the transcriptomic profiles between Mac_Ccl8 and other myeloid cell types, we found that Mac_Ccl8 highly expressed a series of transcription factors, such as Spi1, Maf (c-Maf) and Fli1 (Fig. 3H). Spi1 and Maf are reported to regulate the transcriptional activity of CSF1R, and Maf enhances Ccl8 promoter activity at the transcription level[36–38]. Then, applying single cell trajectory analysis, we found that Mac_Ccl8 was potentially derived from monocytes. Pseudotime analysis revealed later evolution state of Mac_Ccl8 when monocytes were set as the progenitor (Figure 3I, J). Along this trajectory, the expression level of Ccl8 gene rapidly upregulation at the end state, other genes, including Mrc1, Apoe, C1qb, C1qc, and Folr2 also exhibited same expression pattern, whereas expression level of Fn1, Chil3, and Thbs1 genes were higher in early state (Figure 3K).

**Figure 3:**
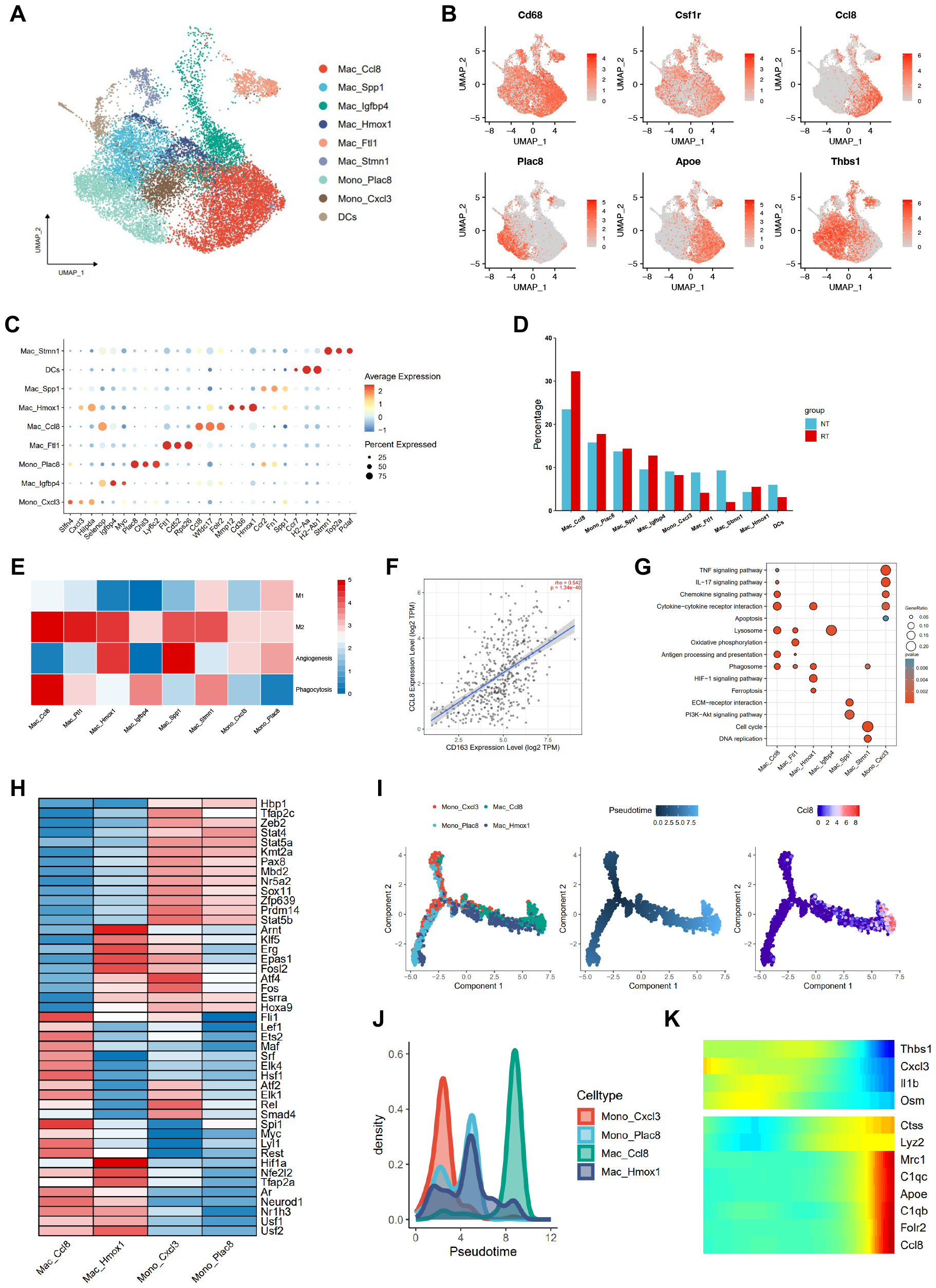
Identification of M2-like Ccl8^high^ Macrophages in the LLC-bearing murine model. (A) UMAP plot showing the annotation of macrophage populations in LLC tumors. (B) Feature plot and (C) Dotplot showing marker genes of macrophage populations. (D) Cell proportion of each cell type in NT and RT groups. (E) Heatmap displaying scores of M1, M2, angiogenesis, phagocytosis for each macrophage population. (F) Correlation analysis of CCL8 and CD163 expression level in TCGA-LUAD datasets (Spearman’s rho value = 0.542, p <0.001). (G) The Gene Ontology (GO) enrichment analysis of each macrophage populations. (H) Heatmap displaying transcription factors activity of the Mac_Ccl8, Mac_Hmox1, Mono_Cxcl3, and Mono_Plac8 populations. (I) Development trajectory of macrophages populations predicted by Monocle2. (J) The cell density and (K) gene expression patterns along with the pseudotime,

In addition, we identified 7 lymphocyte populations based on their transcriptomic files including 2 NK cell populations (NK_Gzma, NK_Gzmd), 4 populations of CD8+ T cell (CD8_Proliferating, CD8_Memory, CD8_Exhausted, and CD8_Ccl2), and a CD4_Treg populations (Figure 4 A, C). The proportion of NK_Gzma, NK_Gzmd and CD8_Exhausted populations was higher in the RT group, while the proportion of CD4_Tregs, CD8_proliferating, CD8_memory populations was higher in the NT group (Figure 4D). Notably, hypofractionated radiotherapy upregulated immune check point genes PD-1, CTLA-4, TIM-3 (Mus musculus Havcr2) in exhausted T cells (Figure 4E, Supplementary Figure 3C). Cell-cell communication analysis revealed that Mac_Ccl8 inhibited CD4 and CD8 T cells activity via immune checkpoint ligand Lgals9 and Cd86 while recruiting NK cells by CCL signaling pathway (Figure 4G). Furthermore, hypofractionated radiotherapy promoted the crosstalk between Mac_Ccl8 and other cell types (Fig 4F, H). For instance, the galectin signaling pathway, including Lgals9 and its receptor Cd44, Cd45, and Havcr2 was significantly upregulated following hypofractionated radiotherapy (Figure 4I).

**Figure 4:**
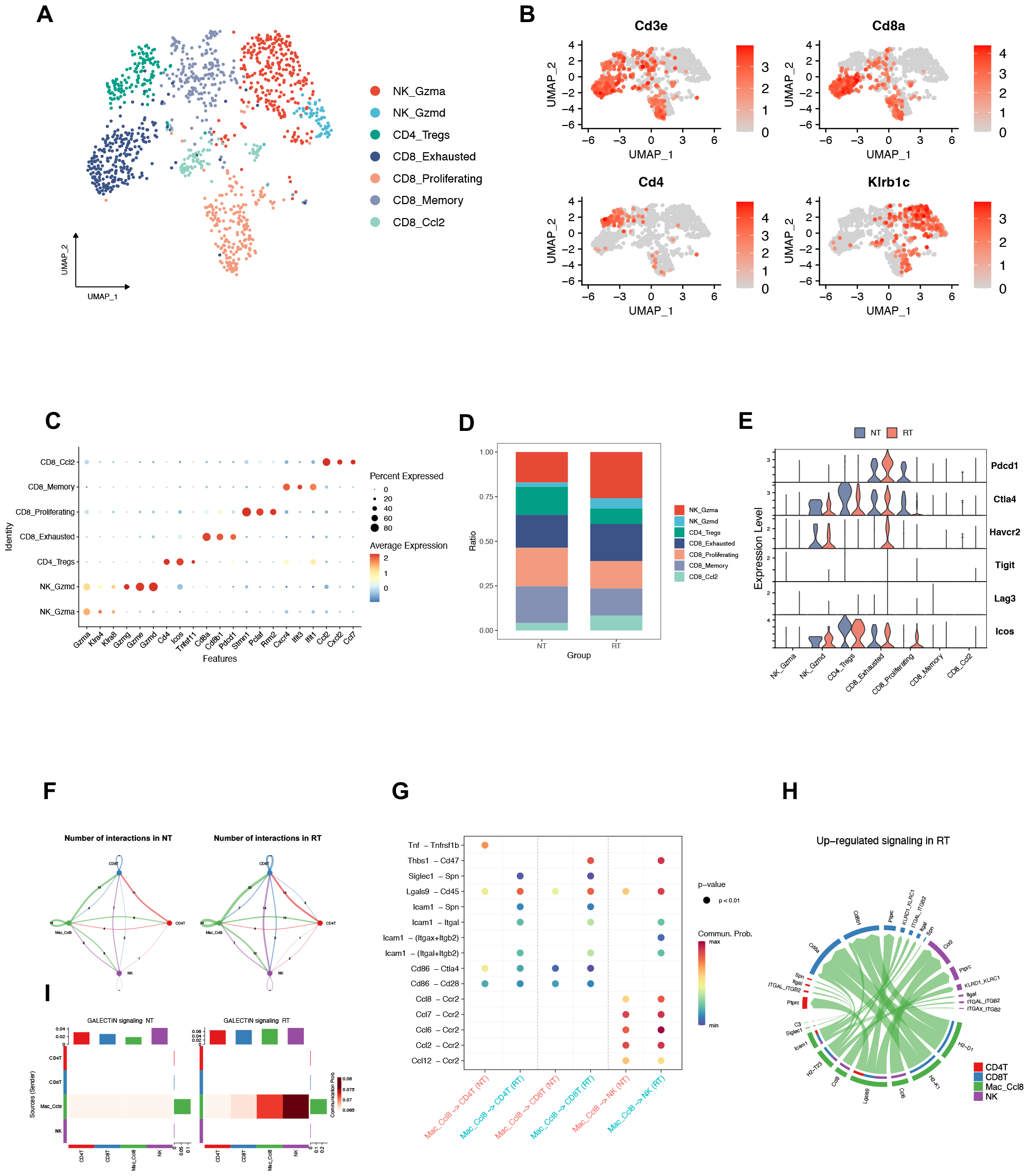
Hypofractionated radiotherapy promoted the crosstalk between the Mac_Ccl8 and lymphocytes. (A) UMAP plot showing the annotation of lymphocyte populations. (B) Feature plot and (C) Dot plot showing marker genes of lymphocyte populations. (D) The proportion of each lymphocyte population in two groups. (E) Violin plots displaying the expression level of immune checkpoint ligand genes in each lymphocyte populations of NT and RT groups. (F) Circle plots showing number of interactions in two groups inferred by CellChat. (G) The comparison of cellular communication probability from Mac_Ccl8 to T and NK cells in between two groups. (H) Chord plot displaying the upregulated signaling pathways in the RT group relative to the NT group. (I) Heatmap of the differential interaction strength of the GALECTIN signaling pathway between two groups.

### Hypofractionated radiotherapy reprograms CCL8^high^ macrophages through the CCL signaling pathway

We sought to investigate whether hypofractionated radiotherapy altered Mac_Ccl8 states. Differentially expressed genes (DEGs) analysis showed that radiation upregulated immune suppressive genes C1qb, Mmp9, and Lgals3bp; pro-phagocytosis gene Icam1; and pro-angiogenesis gene Lyve1, whereas radiation downregulated MHC-II genes H2-Eb1, H2-Aa, and H2-Ab1; pro-inflammatory genes Tnf, Ifnb1, and Il1b (Figure 5A). For chemokines, hypofractionated radiotherapy triggered expression of Ccl8 and Ccl, but downregulated the levels of Ccl3, Ccl4, and Ccl12 (Figure 5B). Additionally, the immunostaining demonstrated an upregulation of CD206 and CCL8 in LLC tumors after radiation (Fig 5C). The GO enrichment analysis of DEGs revealed the enhanced chemokine-activity and chemokine receptor binding, but reduction in antigen presentation of Mac_Ccl8 cluster after treating with hypofractionated radiotherapy (Figure 5D). Gene set enrichment analysis (GSEA) validated downregulation of IFN-γ and TNF signaling in RT Mac_Ccl8, indicating that radiation contribute to its anti-inflammatory function and M2-like polarization (Figure 5E).

**Figure 5:**
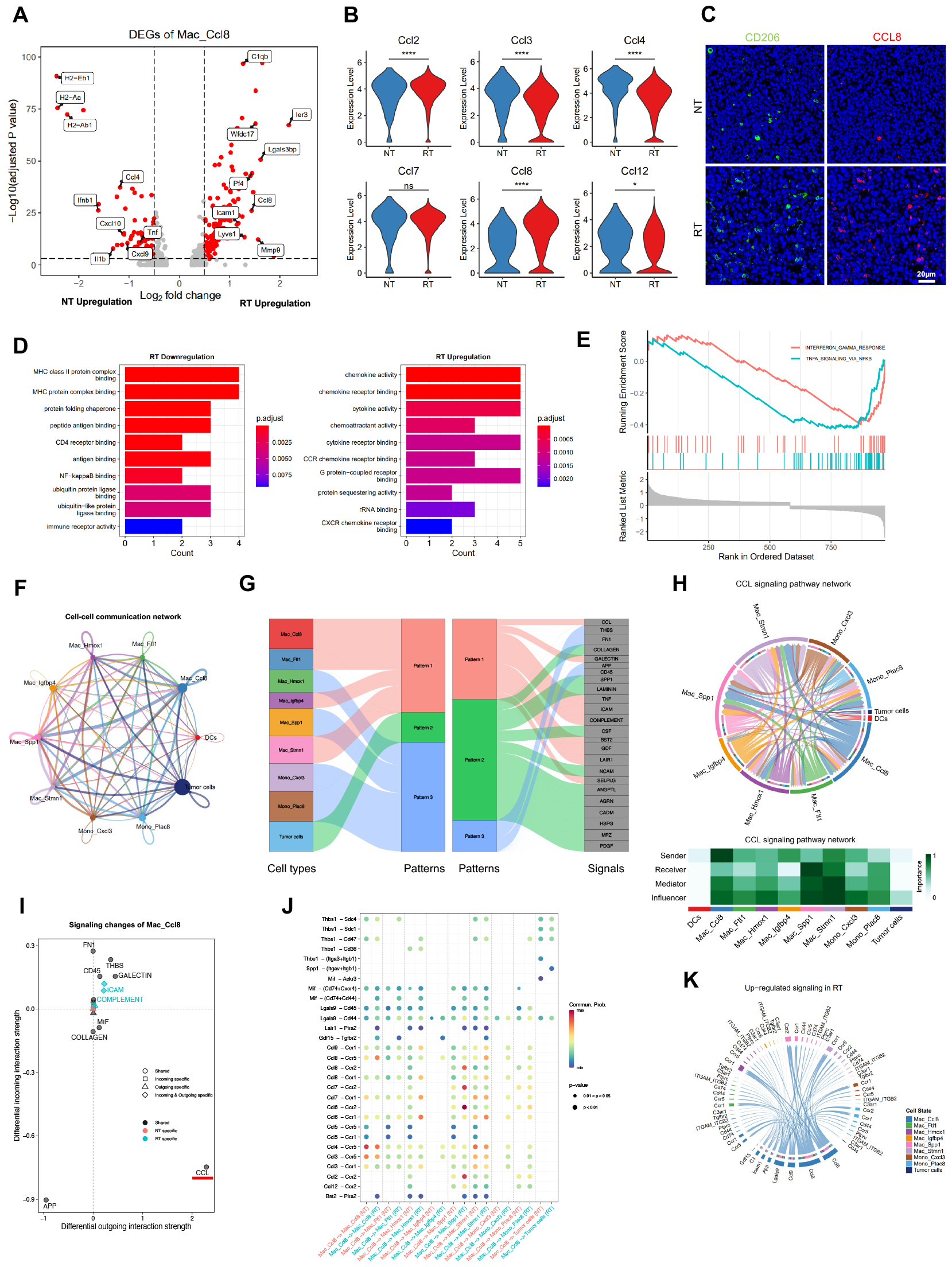
Hypofractionated radiotherapy reprograms CCL8high macrophages through the CCL signaling pathway. (A) Volcano plot showing differentially expressed genes of the Mac_Ccl8 between the NT and RT groups. Adjusted p value <0.05, two-sided Wilcoxon test. (B) Violin plots comparing the expression of Ccl2, Ccl3, Ccl4, Ccl7, Ccl8, and Ccl12 in the NT and RT groups. Unpaired two-sided Wilcoxon test. (C) Representative examples of multiplex immunofluorescent labeling CD206 and CCL8. Green, CD206; Red, CCL8; Blue, DAPI. (D) Bar plots showing the GO enrichment analysis of upregulation and downregulation genes in the RT group. (E) Differences in IFN-Gamma and TNF pathways activity between two groups inferred by GSEA. (F) Cell-cell communication network between myeloid populations and LLC cells. (G) River plot displaying communication patterns of different cell types. (H) Chord plot (top) and heatmap (bottom) showing communication network of CCL signaling pathway in different cell types. (I) Weighted network analysis of differential interaction strength of signals in the Mac_Ccl8 population between two groups. (J) The comparison of cellular communication probability from Mac_Ccl8 to other myeloid populations between two groups. (K) Chord plot displaying the upregulated signaling pathways in the RT group relative to the NT group. *, P < 0.05; **, P < 0.01; ***, P < 0.001; ****, P < 0.0001; ns, not significant.

Next, cell-cell communication network detected 26 signaling pathways between tumor cells and myeloid cells, and that Mac_Ccl8 was the major source communicating each other in macrophage populations (Figure 5F). We then recognized 3 patterns of those signaling pathways. As shown in Figure 5G, Mac_Ccl8 exhibited unique outgoing cellular communication pattern, and the cellular communication pattern 1 featured chemokine signaling pathway (CCL, specifically referring to C-C chemokine in this context). Mac_Ccl8 was identified as the strongest signaling sender whereas Mac_Spp1 was the most significant signaling receiver. Tumor cells were barely involved in cellular communication mediated by CCL signaling pathway (Figure 5H). Furthermore, radiation elevated number and strength of interactions between cells (Supplementary Figure S2G, H). Differential interaction strength analysis demonstrated, as expected, radiation enhanced the communication strength through CCL signaling pathway (Figure 5I). As the major source of CCL signal, radiation augmented communication probability from Mac_Ccl8 to monocyte and macrophage populations including Mac_Ccl8 itself through interaction between CCL8 and CCR receptors (Figure 5J, K). Moreover, we found that recombinant CCL8 protein at concentrations of 5ng/ml and 10ng/ml did not inhibit the viability of LLC cells in vitro. Similarly, adding recombinant CCL8 protein did not affect the colony formation of LLC cells. Thus, Mac_Ccl8 did not promote LLC growth directly (Supplementary Figure S4A, B). Overall, hypofractionated radiotherapy reprogramed CCL8^high^ macrophages toward an immune suppressive state via strengthened CCL signaling-dependent cellular communication between each subset of myeloid cells.

### Hypofractionated radiotherapy promotes M2-like Ccl8^high^ macrophages infiltration and leads to poor prognosis

To investigate whether Mac_Ccl8 in hypofractionated radiotherapy treated s.c. LLC murine model shared similarities with a certain subset in human, we mapped the Ccl8^high^ macrophages signature to well-established myeloid cell atlas -- the Pan-myeloid datasets[32]. Surprisingly, the result from the Pan-myeloid dataset supported that the Mac_Ccl8 signature from our dataset was correlated to the human Macro_C1QC subset, and similarly, human Macro_C1QC was indeed classified as M2 state and exhibited phagocytosis function (Supplementary Figure S2D). Thus, we conducted survival analysis by applying expression profile and clinical data from the TCGA-LUAD. The result suggested that high expression of Mac_Ccl8 gene signatures was associated with worse survival outcomes in LUAD and HNSC patients (Figure 6A).

**Figure 6:**
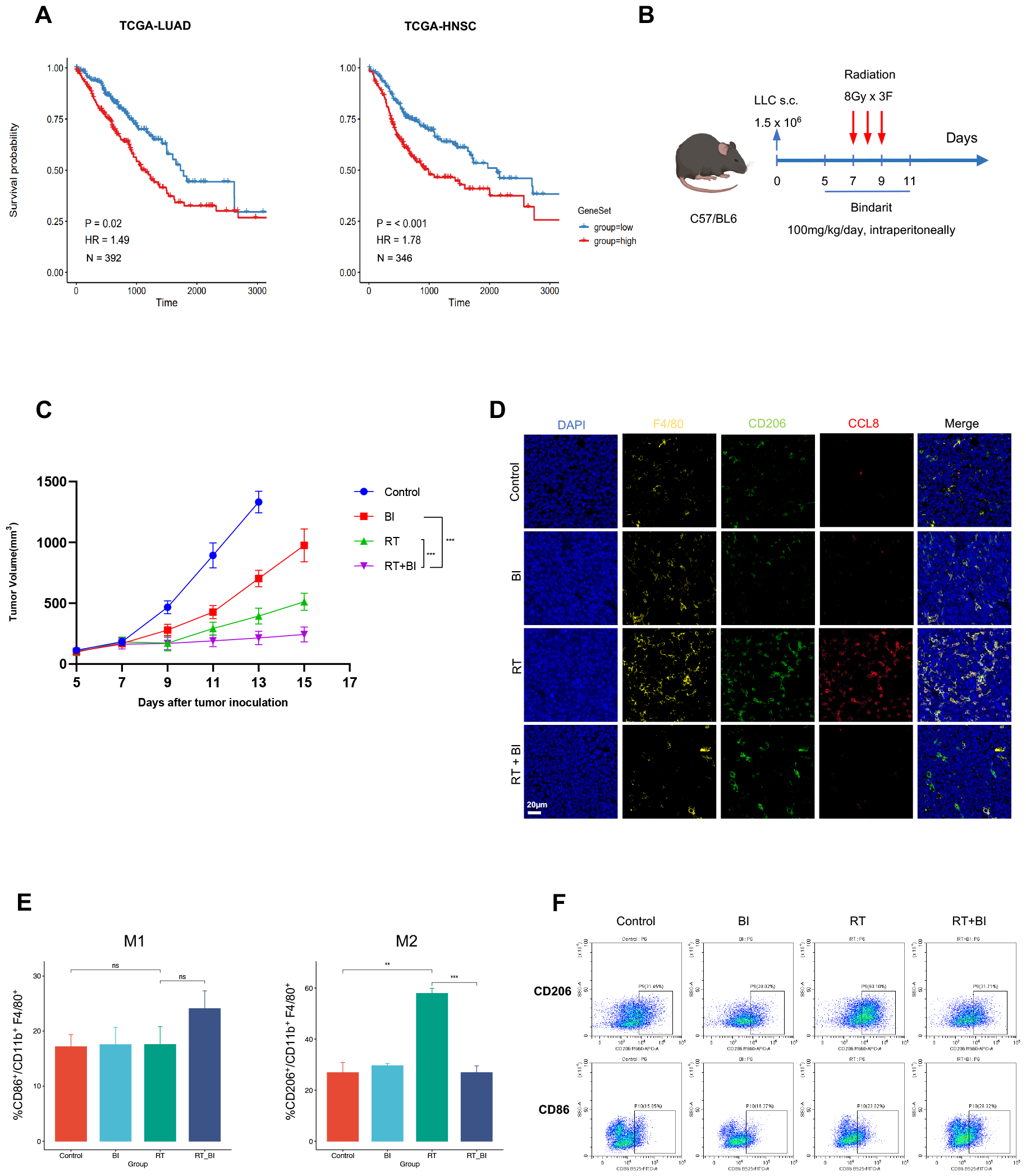
Hypofractionated radiotherapy promotes M2-like Ccl8^high^ macrophages infiltration and leads to poor prognosis. (A) Kaplan-Meier plots showing worse clinical prognosis in the LUAD and HNSC patients with the higher expression level of the Mac_Ccl8 signature. HR, hazard ratio. (B) Schematic diagram of the combination treatment with hypofractionated radiotherapy and the Bindarit. The intraperitoneally administration of Bindarit began from day 5 to day 11 post-tumor injection, and radiation treatment was initiated from day 7 to day 9 post-tumor injection. (C) Growth curves of tumors in LLC-bearing mice in the indicated treatment groups. (D) Representative examples in the indicated treatment groups of multiplex immunofluorescent labeling F4/80, CD206 and CCL8. Yellow, F4/80, Green, CD206; Red, CCL8; Blue, DAPI. (E) Percentages of M1 and M2 macrophages in the specific treatment groups analyzed by flow cytometry. (F) Representative flow cytometry panels showing M2 macrophages (top) and M1 macrophages (bottom). BI, Bindarit; RT, Radiation therapy; RT+BI, the combination therapy of the radiation and the Bindarit.

We further assessed the antitumor effects of the CCL signal (CCL8, CCL7, and CCL2) inhibitor Bindarit in combination with hypofractionated radiotherapy in s.c. LLC murine model. We found that hypofractionated radiotherapy combined with Bindarit significantly prolong the period of local tumor control relative to hypofractionated radiotherapy alone (P<0.001, Figure 6 B, C). By immunostaining, we confirmed a marked influx of CD206+ and CCL8+ macrophages following hypofractionated radiotherapy alone relative to combination treatment, with most pronounced infiltration of M2-like Ccl8^high^ macrophages seen after hypofractionated radiotherapy alone (Figure 6D). We then collected s.c. LLC tumors at early-stage after radiation to evaluated myeloid populations by flow cytometry. The combination therapy reduced the M2 macrophage population and increased the M1 macrophage population as compared to hypofractionated radiotherapy alone (Figure 6E).

## Discussion

In this study, we successfully identified a distinct M2-like population of macrophage with high expression of Ccl8 level at early-stage post-treatment of hypofractionated radiation in the s.c. LLC murine model, which is considered as immunologically cold tumors. Remarkably, hypofractionated radiation not only promoted CCL8^high^ macrophages infiltration but also contributed to M2-like CCL8^high^ macrophages reprogramming, including upregulated immunosuppressive genes (such as C1qb, Mmp9, and Lgals3bp), downregulated antigen-presenting genes (such as H2-Eb1, H2-Aa, and H2-Ab1),and a crosstalk with T cells via immune checkpoint ligands Lgals9 and Cd86, leading to cytotoxic T cell exhaustion and poor prognosis of the patients. Mechanistically, hypofractionated radiation amplified CCL signaling-dependent cellular communication in the tumor immune microenvironment, thus further enhancing pro-tumorigenic functions of the CCL8^high^ macrophage population. Indeed, hypofractionated radiation in combination with Bindarit, a CCL signal inhibitor, reduced M2-like Ccl8^high^ macrophages infiltration and extended the duration of local tumor control in the LLC-bearing mice. Several findings emerged from our analysis that, if further explored, may help to better understand the underlying mechanisms of converting immunologically cold tumors into hot state, and this novel combination treatment strategy potentially develops new vantage points of NSCLC cancer therapy.

TAMs are implicated in promoting tumorigenesis both at primary and metastatic sites. They facilitate tumor cell growth, invasion, angiogenesis and suppress cytotoxic T cells and natural killer (NK) cells, contributing to immune evasion[12–14]. Consequently, TAM infiltration post-radiotherapy is a critical factor in the establishment of an immunosuppressive environment, leading to radiotherapy resistance. Zhang et al.[39] have comprehensively reviewed the mechanisms underlying TAMs recruitment within radiation. Preclinical studies have shown that radiotherapy promotes CCL2, CSF, and HIF-CXCR4 signaling pathways enhancing TAM infiltration. Our findings are consistent with these reports, demonstrating upregulation of the CSF signaling pathway at early-stage following hypofractionated radiotherapy. Notably, our data suggest a significant role for both CCL8 and CCL2 in TAMs recruitment. Leveraging single-cell RNA sequencing, we identified a specific TAM subpopulation, Mac_Ccl8, which is reprogrammed by hypofractionated radiotherapy. Characterized by high Ccl8 expression, Mac_Ccl8 facilitates the recruitment of other macrophages via the CCL signaling pathway. A recent study has highlighted hypofractionated radiotherapy-induced senescence signatures in macrophages, including genes such as Ccl8, Ccl2, Ccl7, Apoe, and Csf1r[40]. The Mac_Ccl8 subpopulation in our study also highly expressed these genes, which relates with worse overall outcomes in NSCLC patients. Collectively, these insights position Mac_Ccl8 as a potential therapeutic target to mitigate the immunosuppressive impact of hypofractionated radiotherapy in tumor microenvironment.

Chemokines are recognized as crucial cytokines in shaping tumor microenvironment through the recruitment of immune cells such as TAMs, MDSCs, and lymphocytes[12,14,41]. Under inflammatory condition, chemokines also induce macrophage polarization towards either M1 or M2 phenotypes[42]. Nevertheless, the mechanisms underlying radiation related immune responses mediated by chemokines remain unclear. Contrary to findings in a glioblastoma research where CCL8 secreted by TAMs was shown to promote tumor growth and invasion[24], our in vitro studies did not demonstrate a similar effect of CCL8 on the growth or invasion of LLC cells. Bindarit is a small molecule inhibitor, which targets CCL signals (CCL8, CCL7, and CCL2) by downregulating the NF-κB pathway[43]. Given that hypofractionated radiotherapy enhanced Ccl8^high^ macrophages infiltration and upregulated CCL signals in a s.c. LLC murine model, we explored a combined therapy of hypofractionated radiotherapy and Bindarit to mitigate TAM infiltration. This combination reduced M2 macrophage infiltration, suggesting that hypofractionated radiotherapy elicits a crosstalk between Ccl8^high^ macrophages and other myeloid cells via the CCL signaling pathway, triggering a cascade effect that recruits and polarizes TAMs. The combination therapy disrupts this cascade effect, thereby prolonging local control of hypofractionated radiotherapy.

Single-cell RNA sequencing offers high resolution transcriptional profiles, yet this study acknowledges three primary limitations. First, while cellular communication analysis indicated an upregulation of the Galectin 9 pathway between Ccl8^high^ macrophages and lymphocytes, the minimal lymphocyte infiltration in LLC tumors constrained our ability to confirm the role of Ccl8^high^ macrophages in T cell exhaustion. Second, the subcutaneous murine tumor model may not fully replicate the tumor microenvironment characteristics observed in orthotopic models. Third, in this study, we only delved on the early alterations in the tumor microenvironment following hypofractionated radiotherapy. Future research examining these changes at various time points post hypofractionated radiotherapy should be considered.

## Conclusions

In summary, we have identified a distinct M2-like population of TAMs which marked by highly expressed Ccl8 gene. hypofractionated radiotherapy reprogrammed Ccl8^high^ macrophages through the upregulation of the CCL signaling pathway which further contributed to TAMs recruitment and polarization. Ccl8^high^ macrophages infiltration increased at early-stage following hypofractionated radiotherapy and related to treatment resistance. The combination therapy of hypofractionated radiotherapy and CCL signals inhibitor mitigated M2 TAMs infiltration and extended local control. These results highlight that targeting TAMs can synergize with hypofractionated radiotherapy.

## Acknowledgements

We thank all the members of the Radiation Oncology Translational Research Group (ROTRG) who participated in this article.

## Formatting of funding sources

This research was supported by grants from the National Natural Science Foundation of China (No. 82373196 to J.D. Sui; No. 82073347 to Y.Z. Wu), Chongqing Science and Health Joint Medical Research Project (No. 2022ZDXM028 to J.D. Sui; No. 2023GGXM002 to YZ Wu), Natural Science Foundation of Chongqing City (No. CSTC2021JXJl0165 to YZ Wu; No. cstc2021jxjl130034 to YZ Wu; No. CSTB2023NSCQ-MSX0709 to JD Sui), Chongqing Talent Plan (No. CQYC20210203119 to YZ Wu).

**Supplementary Figure 1:** Transcriptional characteristics of LLC cells. (A) UMAP plot showing the representative samples in two groups. (B) UMAP plot showing the distribution of tumor cells and immune cells. (C) Heatmap showing the inferCNV analysis results. (D) Violin plots displaying the gene expression level in the CSF and SPP1 signaling pathways across different clusters. (E) Chord plot and (F) heatmap displaying the CSF signaling pathway network.

**Supplementary Figure 2:** Transcriptional characteristics of myeloid cells. (A) Differentially expressed genes of each myeloid populations. Red dots represent upregulated genes, Blue dots represent downregulated genes. (B) Violin plots showing the gene expression level of the M1 and M2 signatures in the NT and the RT groups. (C) Dot plot showing the KEGG analysis results of macrophages and monocytes. (D) Representative of the Mac_Cc8 signature enrichment in the panmyeloid database (http://panmyeloid.cancer-pku.cn/). (D) Dot plot showing cellular communication signaling pathways between Mac_Ccl8 and other cell types. (F) Hierarchy plot showting the CCL signaling pathway network between myeloid cell populations. (G) Bar plot and (H) heatmap showing the differential interaction number and strength between the NT and the RT groups.

**Supplementary Figure 3:** Cell-cell communication analysis between Mac_Ccl8 and lymphocytes. (A) Dot plot showing the KEGG analysis results of different lymphocytes. (B) Dot plot showing the signaling pathways between Mac_Ccl8 and lymphocytes. (C) Chord plots showing the PD-L1, CD86, and GALECTIN signaling pathway networks. (D) Strength of interactions in the NT and RT groups. (E) Heatmap showing differential communication strength of MHC-I signaling between the NT and the RT groups.

**Supplementary Figure 4:** Recombinant CCL8 protein did not promote LLC cells proliferation in vitro. (A) Cell Viability in the CCK-8 assay in the indicated CCL8 protein concentration group. (B) Colony formation assay for LLC cells with the addition of different concentrations of CCL8 protein concentrations. (C) The gating strategy for isolating macrophages. Live cells were gated first. Then M1 macrophages were gated on CD45^+^CD11b^+^F4/80^+^CD86^+^ population. M2 macrophages were gated on CD45^+^CD11b^+^F4/80^+^CD206^+^ poplulation.

## Supplementary Materials

Figure S1: Transcriptional characteristics of LLC cells

Figure S2: Tran-scriptional characteristics of myeloid cells

Figure S3: Cell-cell communication analysis between Mac_Ccl8 and lymphocytes

Figure S4: Recombinant CCL8 protein did not promote LLC cells proliferation in vitro

Table S1: the list of gene signatures

### Research data

We are currently in the process of uploading our single-cell RNA sequencing data to the GEO (Gene Expression Omnibus) repository. This task will be completed prior to the acceptance of our work.

## References

1. Vinod, S.K.; Hau, E. Radiotherapy Treatment for Lung Cancer: Current Status and Future Directions. Respirology 2020, 25, 61–71, doi:10.1111/resp.13870.

2. Cheng, M.; Jolly, S.; Quarshie, W.O.; Kapadia, N.; Vigneau, F.D.; Kong, F.-M. (Spring) Modern Radiation Further Improves Survival in Non-Small Cell Lung Cancer: An Analysis of 288,670 Patients. J. Cancer 2019, 10, 168–177, doi:10.7150/jca.26600.

3. Chang, J.Y.; Mehran, R.J.; Feng, L.; Verma, V.; Liao, Z.; Welsh, J.W.; Lin, S.H.; O’Reilly, M.S.; Jeter, M.D.; Balter, P.A.; et al. Stereotactic Ablative Radiotherapy for Operable Stage I Non-Small-Cell Lung Cancer (Revised STARS): Long-Term Results of a Single-Arm, Prospective Trial with Prespecified Comparison to Surgery. Lancet Oncol. 2021, 22, 1448–1457, doi:10.1016/S1470-2045(21)00401-0.

4. Chang, J.Y.; Lin, S.H.; Dong, W.; Liao, Z.; Gandhi, S.J.; Gay, C.M.; Zhang, J.; Chun, S.G.; Elamin, Y.Y.; Fossella, F.V.; et al. Stereotactic Ablative Radiotherapy with or without Immunotherapy for Early-Stage or Isolated Lung Parenchymal Recurrent Node-Negative Non-Small-Cell Lung Cancer: An Open-Label, Randomised, Phase 2 Trial. The Lancet 2023, 402, 871–881, doi:10.1016/S0140-6736(23)01384-3.

5. Zhang, Z.; Liu, X.; Chen, D.; Yu, J. Radiotherapy Combined with Immunotherapy: The Dawn of Cancer Treatment. Signal Transduct. Target. Ther. 2022, 7, 258, doi:10.1038/s41392-022-01102-y.

6. Herrera, F.G.; Ronet, C.; Ochoa de Olza, M.; Barras, D.; Crespo, I.; Andreatta, M.; Corria-Osorio, J.; Spill, A.; Benedetti, F.; Genolet, R.; et al. Low Dose Radiotherapy Reverses Tumor Immune Desertification and Resistance to Immunotherapy. Cancer Discov 2021, doi:10.1158/2159-8290.CD-21-0003.

7. Zhu, M.; Yang, M.; Zhang, J.; Yin, Y.; Fan, X.; Zhang, Y.; Qin, S.; Zhang, H.; Yu, F. Immunogenic Cell Death Induction by Ionizing Radiation. Front. Immunol. 2021, 12, 705361, doi:10.3389/fimmu.2021.705361.

8. Deng, L.; Liang, H.; Xu, M.; Yang, X.; Burnette, B.; Arina, A.; Li, X.-D.; Mauceri, H.; Beckett, M.; Darga, T.; et al. STING-Dependent Cytosolic DNA Sensing Promotes Radiation-Induced Type I Interferon-Dependent Antitumor Immunity in Immunogenic Tumors. Immunity 2014, 41, 843–852, doi:10.1016/j.immuni.2014.10.019.

9. Woods, D.M.; Ramakrishnan, R.; Laino, A.S.; Berglund, A.; Walton, K.; Betts, B.C.; Weber, J.S. Decreased Suppression and Increased Phosphorylated STAT3 in Regulatory T Cells Are Associated with Benefit from Adjuvant PD-1 Blockade in Resected Metastatic Melanoma. Clin. Cancer Res. Off. J. Am. Assoc. Cancer Res. 2018, 24, 6236–6247, doi:10.1158/1078-0432.CCR-18-1100.

10. Oweida, A.J.; Darragh, L.; Phan, A.; Binder, D.; Bhatia, S.; Mueller, A.; Court, B.V.; Milner, D.; Raben, D.; Woessner, R.; et al. STAT3 Modulation of Regulatory T Cells in Response to Radiation Therapy in Head and Neck Cancer. JNCI J. Natl. Cancer Inst. 2019, 111, 1339–1349, doi:10.1093/jnci/djz036.

11. Boutilier, A.J.; Elsawa, S.F. Macrophage Polarization States in the Tumor Microenvironment. Int. J. Mol. Sci. 2021, 22, 6995, doi:10.3390/ijms22136995.

12. Mantovani, A.; Allavena, P.; Marchesi, F.; Garlanda, C. Macrophages as Tools and Targets in Cancer Therapy. Nat. Rev. Drug Discov. 2022, 21, 799–820, doi:10.1038/s41573-022-00520-5.

13. Chen, Y.; Song, Y.; Du, W.; Gong, L.; Chang, H.; Zou, Z. Tumor-Associated Macrophages: An Accomplice in Solid Tumor Progression. J. Biomed. Sci. 2019, 26, 78, doi:10.1186/s12929-019-0568-z.

14. Cassetta, L.; Pollard, J.W. A Timeline of Tumour-Associated Macrophage Biology. Nat. Rev. Cancer 2023, 23, 238–257, doi:10.1038/s41568-022-00547-1.

15. Bule, P.; Aguiar, S.I.; Aires-Da-Silva, F.; Dias, J.N.R. Chemokine-Directed Tumor Microenvironment Modulation in Cancer Immunotherapy. Int. J. Mol. Sci. 2021, 22, 9804, doi:10.3390/ijms22189804.

16. Nagarsheth, N.; Wicha, M.S.; Zou, W. Chemokines in the Cancer Microenvironment and Their Relevance in Cancer Immunotherapy. Nat. Rev. Immunol. 2017, 17, 559–572, doi:10.1038/nri.2017.49.

17. Wang, L.; Jiang, J.; Chen, Y.; Jia, Q.; Chu, Q. The Roles of CC Chemokines in Response to Radiation. Radiat. Oncol. 2022, 17, 63, doi:10.1186/s13014-022-02038-x.

18. Liang, H.; Deng, L.; Hou, Y.; Meng, X.; Huang, X.; Rao, E.; Zheng, W.; Mauceri, H.; Mack, M.; Xu, M.; et al. Host STING-Dependent MDSC Mobilization Drives Extrinsic Radiation Resistance. Nat. Commun. 2017, 8, 1736, doi:10.1038/s41467-017-01566-5.

19. Kalbasi, A.; Komar, C.; Tooker, G.M.; Liu, M.; Lee, J.W.; Gladney, W.L.; Ben-Josef, E.; Beatty, G.L. Tumor-Derived CCL2 Mediates Resistance to Radiotherapy in Pancreatic Ductal Adenocarcinoma. Clin. Cancer Res. Off. J. Am. Assoc. Cancer Res. 2017, 23, 137–148, doi:10.1158/1078-0432.CCR-16-0870.

20. Wang, X.; Yang, X.; Tsai, Y.; Yang, L.; Chuang, K.-H.; Keng, P.C.; Lee, S.O.; Chen, Y. IL-6 Mediates Macrophage Infiltration after Irradiation via Up-Regulation of CCL2/CCL5 in Non-Small Cell Lung Cancer. Radiat. Res. 2017, 187, 50–59, doi:10.1667/RR14503.1.

21. Dangaj, D.; Bruand, M.; Grimm, A.J.; Ronet, C.; Barras, D.; Duttagupta, P.A.; Lanitis, E.; Duraiswamy, J.; Tanyi, J.L.; Benencia, F.; et al. Cooperation between Constitutive and Inducible Chemokines Enables T Cell Engraftment and Immune Attack in Solid Tumors. Cancer Cell 2019, 35, 885–900.e10, doi:10.1016/j.ccell.2019.05.004.

22. Liu, X.; Zeng, L.; Zhou, Y.; Zhao, X.; Zhu, L.; Zhang, J.; Pan, Y.; Shao, C.; Fu, J. P21 Facilitates Macrophage Chemotaxis by Promoting CCL7 in the Lung Epithelial Cell Lines Treated with Radiation and Bleomycin. J. Transl. Med. 2023, 21, 1–19, doi:10.1186/s12967-023-04177-5.

23. Korbecki, J.; Kojder, K.; Simińska, D.; Bohatyrewicz, R.; Gutowska, I.; Chlubek, D.; Baranowska-Bosiacka, I. CC Chemokines in a Tumor: A Review of Pro-Cancer and Anti-Cancer Properties of the Ligands of Receptors CCR1, CCR2, CCR3, and CCR4. Int. J. Mol. Sci. 2020, 21, 8412, doi:10.3390/ijms21218412.

24. Zhang, X.; Chen, L.; Dang, W.; Cao, M.; Xiao, J.; Lv, S.; Jiang, W.; Yao, X.; Lu, H.; Miao, J.; et al. CCL8 Secreted by Tumor-Associated Macrophages Promotes Invasion and Stemness of Glioblastoma Cells via ERK1/2 Signaling. Lab. Invest. 2020, 100, 619–629, doi:10.1038/s41374-019-0345-3.

25. Hao, Y.; Hao, S.; Andersen-Nissen, E.; Mauck, W.M.; Zheng, S.; Butler, A.; Lee, M.J.; Wilk, A.J.; Darby, C.; Zager, M.; et al. Integrated Analysis of Multimodal Single-Cell Data. Cell 2021, 184, 3573–3587.e29, doi:10.1016/j.cell.2021.04.048.

26. Korsunsky, I.; Millard, N.; Fan, J.; Slowikowski, K.; Zhang, F.; Wei, K.; Baglaenko, Y.; Brenner, M.; Loh, P.; Raychaudhuri, S. Fast, Sensitive and Accurate Integration of Single-Cell Data with Harmony. Nat. Methods 2019, 16, 1289–1296, doi:10.1038/s41592-019-0619-0.

27. Garcia-Alonso, L.; Holland, C.H.; Ibrahim, M.M.; Turei, D.; Saez-Rodriguez, J. Benchmark and Integration of Resources for the Estimation of Human Transcription Factor Activities. Genome Res. 2019, 29, 1363–1375, doi:10.1101/gr.240663.118.

28. Wu, T.; Hu, E.; Xu, S.; Chen, M.; Guo, P.; Dai, Z.; Feng, T.; Zhou, L.; Tang, W.; Zhan, L.; et al. clusterProfiler 4.0: A Universal Enrichment Tool for Interpreting Omics Data. The Innovation 2021, 2, doi:10.1016/j.xinn.2021.100141.

29. Hänzelmann, S.; Castelo, R.; Guinney, J. GSVA: Gene Set Variation Analysis for Microarray and RNA-Seq Data. BMC Bioinformatics 2013, 14, 7, doi:10.1186/1471-2105-14-7.

30. Qiu, X.; Mao, Q.; Tang, Y.; Wang, L.; Chawla, R.; Pliner, H.A.; Trapnell, C. Reversed Graph Embedding Resolves Complex Single-Cell Trajectories. Nat. Methods 2017, 14, 979–982, doi:10.1038/nmeth.4402.

31. Jin, S.; Guerrero-Juarez, C.F.; Zhang, L.; Chang, I.; Ramos, R.; Kuan, C.-H.; Myung, P.; Plikus, M.V.; Nie, Q. Inference and Analysis of Cell-Cell Communication Using CellChat. Nat. Commun. 2021, 12, 1088, doi:10.1038/s41467-021-21246-9.

32. Cheng, S.; Li, Z.; Gao, R.; Xing, B.; Gao, Y.; Yang, Y.; Qin, S.; Zhang, L.; Ouyang, H.; Du, P.; et al. A Pan-Cancer Single-Cell Transcriptional Atlas of Tumor Infiltrating Myeloid Cells. Cell 2021, 184, 792–809.e23, doi:10.1016/j.cell.2021.01.010.

33. Barata, T.; Duarte, I.; Futschik, M.E. Integration of Stemness Gene Signatures Reveals Core Functional Modules of Stem Cells and Potential Novel Stemness Genes. Genes 2023, 14, 745, doi:10.3390/genes14030745.

34. Gulati, G.S.; Sikandar, S.S.; Wesche, D.J.; Manjunath, A.; Bharadwaj, A.; Berger, M.J.; Ilagan, F.; Kuo, A.H.; Hsieh, R.W.; Cai, S.; et al. Single-Cell Transcriptional Diversity Is a Hallmark of Developmental Potential. Science 2020, 367, 405–411, doi:10.1126/science.aax0249.

35. Stanley, E.R.; Chitu, V. CSF-1 Receptor Signaling in Myeloid Cells. Cold Spring Harb. Perspect. Biol.2014, 6, a021857, doi:10.1101/cshperspect.a021857.

36. Aikawa, Y.; Katsumoto, T.; Zhang, P.; Shima, H.; Shino, M.; Terui, K.; Ito, E.; Ohno, H.; Stanley, E.R.; Singh, H.; et al. PU.1-Mediated Upregulation of CSF1R Is Crucial for Leukemia Stem Cell Potential Induced by MOZ-TIF2. Nat. Med. 2010, 16, 580–585, 1p following 585, doi:10.1038/nm.2122.

37. Kikuchi, K.; Iida, M.; Ikeda, N.; Moriyama, S.; Hamada, M.; Takahashi, S.; Kitamura, H.; Watanabe, T.; Hasegawa, Y.; Hase, K.; et al. Macrophages Switch Their Phenotype by Regulating Maf Expression during Different Phases of Inflammation. J. Immunol. 2018, 201, 635–651, doi:10.4049/jimmunol.1800040.

38. Liu, M.; Tong, Z.; Ding, C.; Luo, F.; Wu, S.; Wu, C.; Albeituni, S.; He, L.; Hu, X.; Tieri, D.; et al. Transcription Factor C-Maf Is a Checkpoint That Programs Macrophages in Lung Cancer. J. Clin. Invest. 2020, 130, 2081–2096, doi:10.1172/JCI131335.

39. Zhang, Z.; Liu, X.; Chen, D.; Yu, J. Radiotherapy Combined with Immunotherapy: The Dawn of Cancer Treatment. Signal Transduct. Target. Ther. 2022, 7, 258, doi:10.1038/s41392-022-01102-y.

40. Wu, F.; Zhang, Z.; Wang, M.; Ma, Y.; Verma, V.; Xiao, C.; Zhong, T.; Chen, X.; Wu, M.; Yu, J.; et al. Cellular Atlas of Senescent Lineages in Radiation-or Immunotherapy-Induced Lung Injury by Single-Cell RNA-Sequencing Analysis. Int. J. Radiat. Oncol. Biol. Phys. 2023, S0360-3016(23)00148-7, doi:10.1016/j.ijrobp.2023.02.005.

41. Drouillard, D.; Craig, B.T.; Dwinell, M.B. Physiology of Chemokines in the Cancer Microenvironment. Am. J. Physiol.-Cell Physiol. 2023, doi:10.1152/ajpcell.00151.2022.

42. Ruytinx, P.; Proost, P.; Van Damme, J.; Struyf, S. Chemokine-Induced Macrophage Polarization in Inflammatory Conditions. Front. Immunol. 2018, 9, 1930, doi:10.3389/fimmu.2018.01930.

43. Mora, E.; Guglielmotti, A.; Biondi, G.; Sassone-Corsi, P. Bindarit: An Anti-Inflammatory Small Molecule That Modulates the NFkB Pathway. Cell Cycle 2012, 11, 159–169, doi:10.4161/cc.11.1.18559.

