## Supplementary figures and images for "Single-cell RNA sequencing reveals recruitment of the M2-like CCL8^high^ macrophages in Lewis lung carcinoma-bearing mice following hypofractionated radiotherapy"

### Supplemental Figures

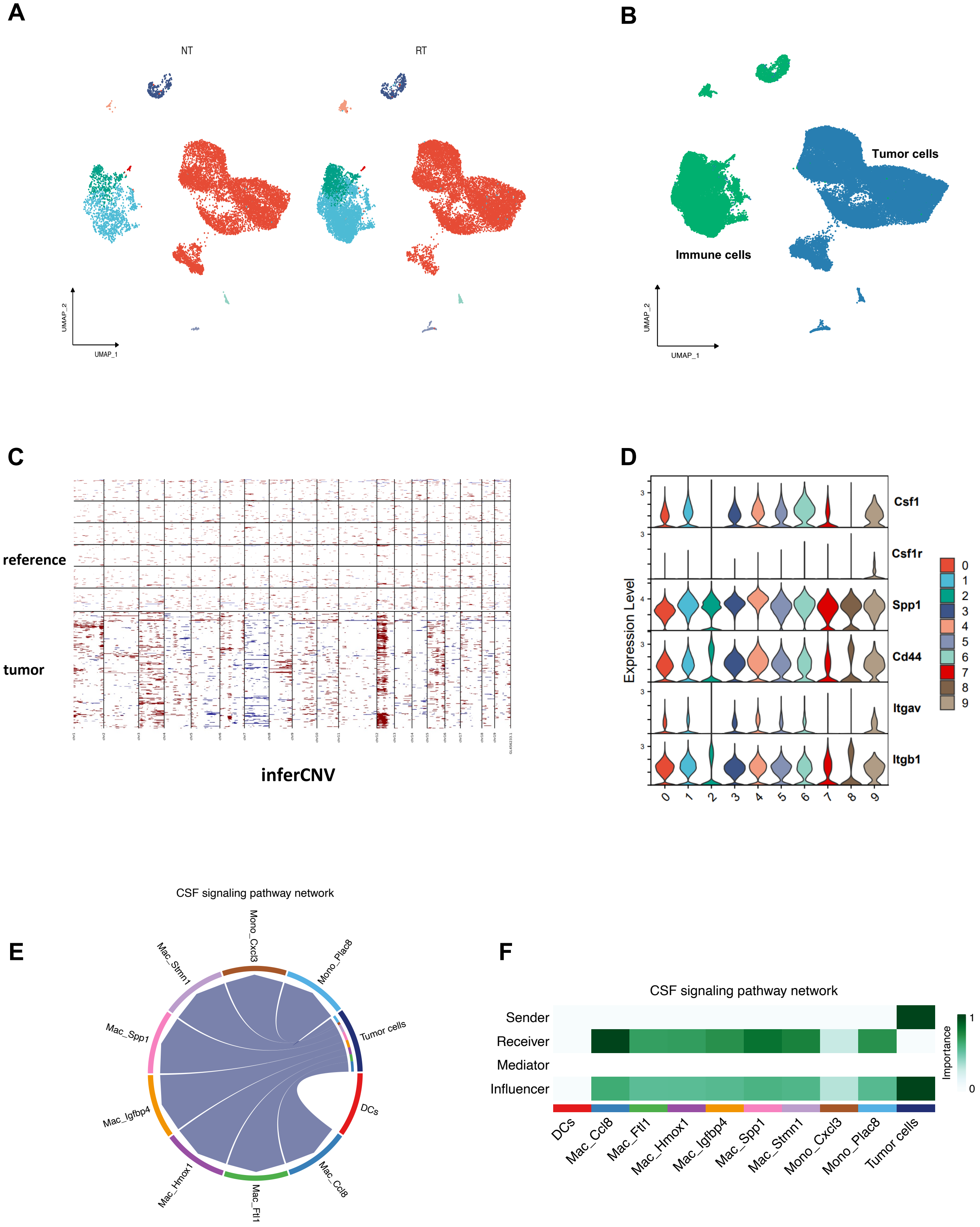

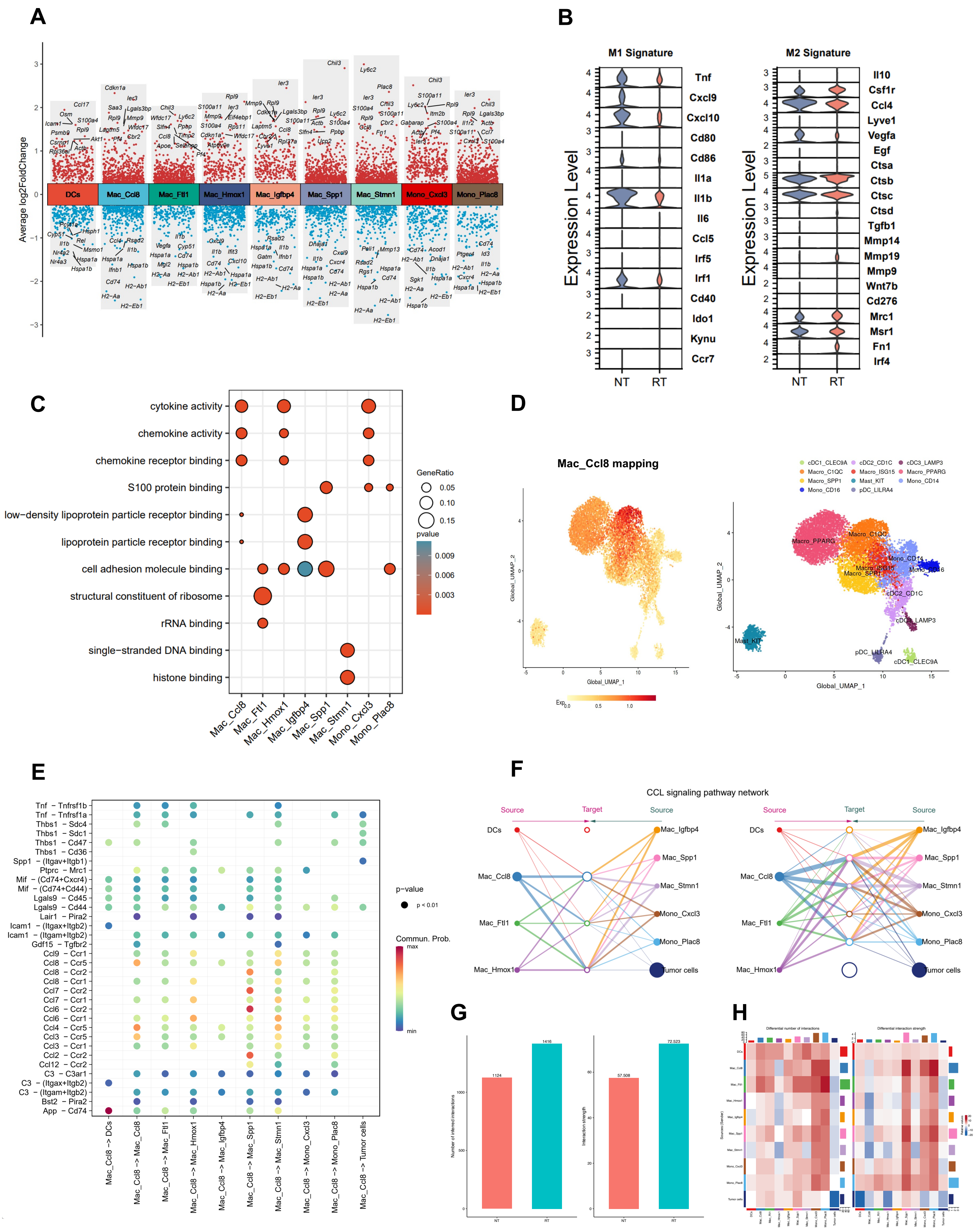

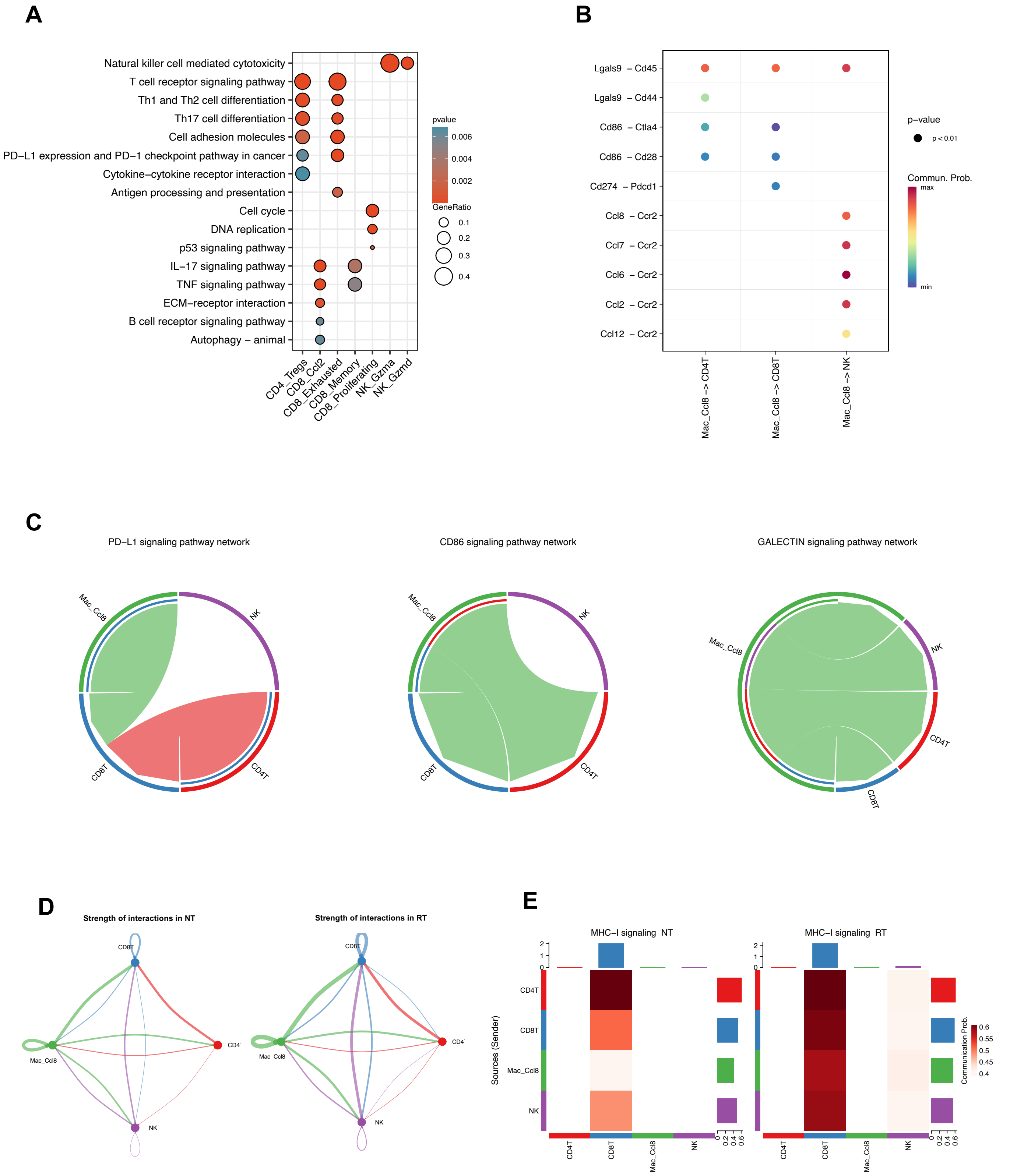

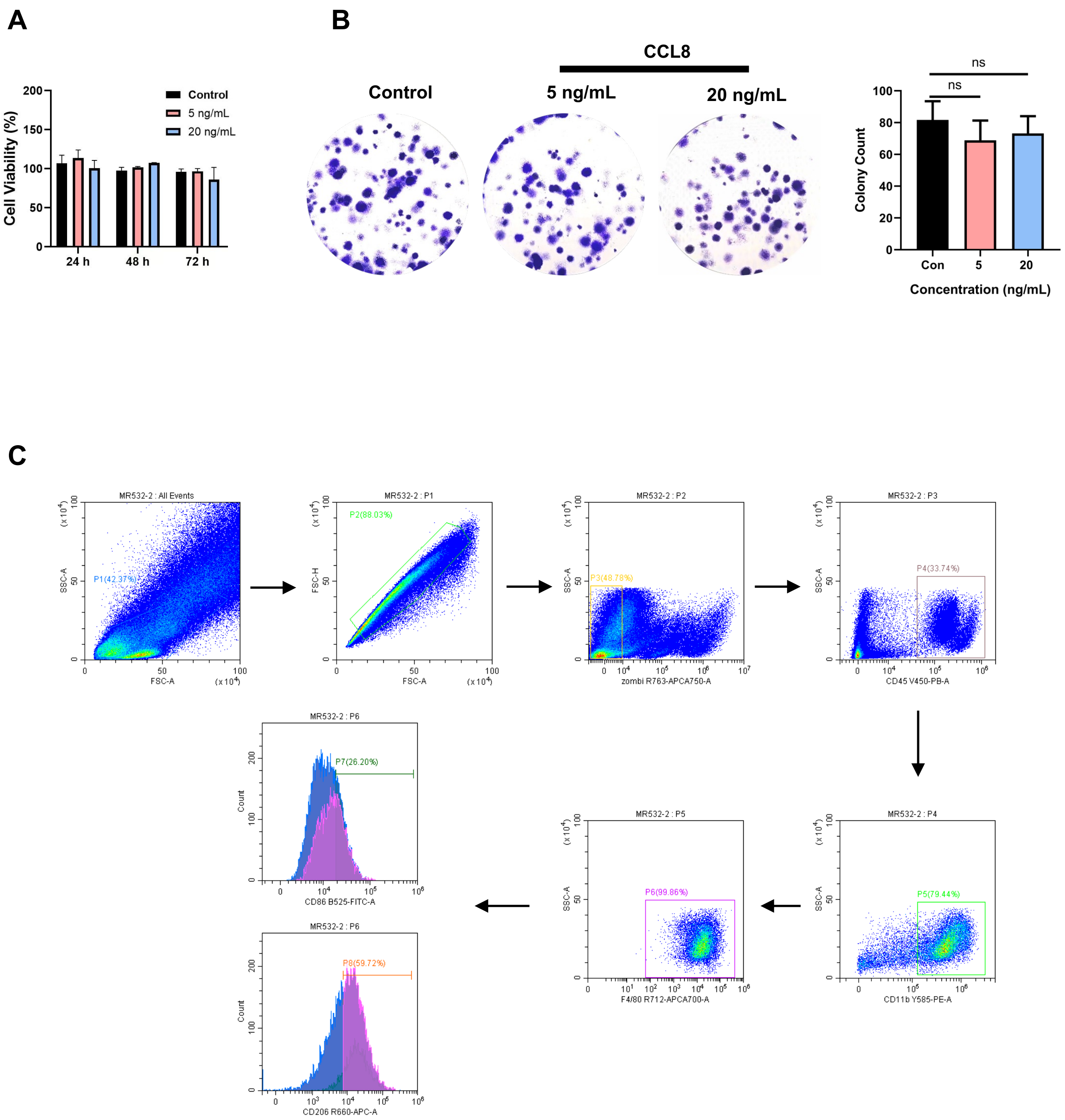
